# Viscoelasticity and Noise Properties Reveal the Formation of Biomemory in Cells

**DOI:** 10.1101/2021.08.06.455392

**Authors:** Evangelos Bakalis, Vassilios Gavriil, Alkiviadis-Constantinos Cefalas, Zoe Kollia, Francesco Zerbetto, Evangelia Sarantopoulou

## Abstract

Living cells are neither perfectly elastic nor liquid and return a viscoelastic response to external stimuli. Nanoindentation provides force distance curves allowing the investigation of cell mechanical properties, and yet, these curves can differ from point to point on cell surface revealing its inhomogeneous character. In the present work, we propose a mathematical method to estimate both viscoelastic and noise properties of cells, as these are depicted on the values of the scaling exponents of relaxation function and power spectral density respectively. The method uses as input the time derivative of the response force in a nanoindentation experiment. Generalized moments method and/or rescaled range analysis are used to study the resulting time series depending on their non-stationary or stationary nature. We conducted experiments in living *Ulocladium Chartarum* spores. We found that spores, in the approaching phase present a viscoelastic behavior with the corresponding scaling exponent in the range 0.25-0.52, and in the retracting phase present a liquid-like behavior with exponents in the range 0.67-0.85. This substantial difference of the scaling exponents in the two phases suggests the formation of biomemory as response of the spores to the indenting AFM mechanical stimulus. The retracting phase may be described as a process driven by bluish noises, while the approaching one is driven by persistent noise.

## Introduction

Living cells are continuously subject to mechanical forces both by surrounding cells and by the microenvironment they belong to. They adapt their bio-response to extracellular environmental conditions by tracing maximum viability lines. ^1,2^ On a single cell, mechanical forces may cause shear, stress, and torsion. The generally accepted scenario is that in a cell the response to mechanical deformations is due to the activation of external cell wall protein-like mechanosensors, which are connected internally with an extended plasma membrane contractile network formed mainly by actin filaments. ^3–6^ Response to external stimuli determines elastic properties of a cell surface and relates them to concrete tasks, for example, softening supports cell mobility and migration. ^3^ Cell elasticity is a measure whose changes are used as indicators for cytotoxicity, malignancy, viability, bio-memory and abnormalities. ^1–9^ Furthermore, changes of cell elasticity, resulting from external stress, have been associated with cell abnormalities such as cancer, cardiomyopathies, generation of diverse dysmorphic phenotypes. ^8,10,11^ This property has been used for on-the-fly cell mechanical phenotyping. ^12^ Cell mechanics, which examines the response of a cell to stimuli of biochemical chemical or physical nature,^13,14^ has been investigated by applying different techniques, including Atomic Force Microscopy nano-indentation (AFM-NI), see a recent comparison of methods to assess cell mechanical properties. ^15^

AFM-NI has been widely used to characterize mechanical properties of both cells and tissues. ^13,16,17^ It uses a tip of well-defined geometry to punch into the cell placed on a solid support, and is commonly used to quantify mechanical properties at sub-cellular resolution. It can also perform precise force measurements at desired cellular locations where the tip of the cantilever is used as indenter. The measured force, attractive or repulsive, corresponds to the interaction between tip atoms with those that belong to the sample surface. The vertical displacement of a cantilever and its deflection are recorded simultaneously and then converted to force distance curves (FDC). FDC’s are registered in both phases, approach (tip moving towards the sample) and retraction (tip withdrawing from the sample).

To a first approximation, the Young’s modulus is estimated by using the Hertz model,^18,19^ which describes the response of an isotropic and fully elastic material under a load, to fit the FDC’s. In this approximation the Young’s modulus is time independent. For complex materials like cells, the estimate of Young’s modulus based solely on the Hertz model and without considering viscoelastic effects (combination of elastic and liquid behaviour) is highly questionable. The Young’s modulus is a time-dependent measure and its estimate is affected both by the thickness of the cell and by the solid support where it is placed on.^20,21^ Experimentally, the non-identical approach and retraction parts of FDC’s are signs of viscoelasticity. A reason for the differences likely is the diverse hydrodynamic drag on the cantilever. Furthermore, in the contact regime, a difference between approach and retraction is an indication of plastic deformations or, most typically, a viscoelastic behaviour of the sample, which underlines the formation of a type of memory (biomemory) because the cell alters its local environment via excessive protein activity at the activation site. ^2^

Cells are not a simple fluid-filled envelope, they contain different active intracellular structures that may display distinct mechanical properties. ^22^ Cells display both solid-like elastic and fluid-like viscous properties, and under external stress typically return a viscoelastic behaviour, which is reflected on a power-law form satisfied by both the creep and the stress relaxation functions. ^23,24^ A vast number of studies based on a variety of techniques showed that the rheological properties of cells are better described by a power law relaxation function of the form *E*(*t*) = *E*_0_(*t/t*_0_)^−*β*^ with 0 ≥ *β* ≥ 1. *E*_0_ is the Young’s modulus at time *t*_0_, which can be chosen arbitrarily and is usually set to 1 sec.^25^ *E*_0_ deviates from the Young’s moduli provided by elastic models which are constant in time. The scaling exponent *β* characterizes the degree of fluidity and energy dissipation upon deformation. A value of *β* = 0 stands for a perfectly elastic solid and a value of *β* = 1 for a Newtonian liquid. Any value of the scaling exponent between these two limits describes a viscoelastic medium. A typical value of *β* for cells lies in the range 0.1-0.3 classifying thus a cell as a viscoelastic solid. ^25–28^ For cells, it has been reported that the dependency of elastic modulus on probing frequency follows a weak power law resulted in the absence of discrete relaxation times in the system. ^29^ For viscoelastic materials, such as cells, their response is not only a function of the instantaneous deformations caused by the exerted mechanical forces but also depends on the history of deformations.^30,31^

In specific fungal and carcinogenic cells, an external stress is always followed by a bio-response for maximum viability via a biomemory cell system. ^1,2,8,9^ A memory kernel can describe the history of deformations, and its form indicates how strong is the memory formed under the action of mechanical forces. For example, a Dirac delta memory kernel describes memory-less deformations, an exponential decay can describe a Poisson distribution of deformations, and a power-law memory kernel accounts for strong memory effects. History dependent deformations in viscoelastic systems, where elastic and viscous properties coexist to varying degrees, may cause non-local effects both in time and space, ^32^ and may be modeled by fractional calculus, ^33^ which is an appropriate framework to model complexity. ^34,35^ Additionally, complexity may be modeled by a fractional Langevin equation where the overall noise may behave as a multiplicative process. The role of such a class of noises has been studied for a variety of systems ranging from ecology, ^36^ to pattern formation, ^37^ and to stability of biological systems. ^38^ Experimentally, the complexity of the mechanical behaviours to deformations in single cells and/or tissues has been pointed out, ^31,39–42^ see also a recent review.^43^

Considering a cell as an incompressible material, its response to a mechanical load may be expressed as a function of the indentation depth and the creep relaxation function through their convolution. ^43^ In an AFM-NI experiment, the response forces form a data set with hierarchical time distribution and define the observation window. The analysis of events in such a window, which usually contains few data points, can infer past and future events only if the process is deterministic or periodic or stationary. For non-stationary processes, like approach and retraction parts of a FDC, one can use more sophisticated methods, appropriate for time series analysis. Among them ^44–49^ the generalized moments method (GMM) is generally one of the more robust and works well even for short time series.^50^ It has been successfully applied in numerous fields, ^51–56^ and it works for non-stationary time series. ^55,56^ For the stationary ones, rescaled range analysis (RA)^57,58^ or some variations thereof ^59^ are the proper analysis methods. Both methods, GMM and RA, deliver the scaling exponent, which is called Hurst exponent, of a stochastic process. Additionally, there is a link between these scaling exponents with the scaling of the power spectral density (PSD) whose value classifies the color of the stochastic process. ^60^ To distinguish the method applied for analysis the symbols with subscripts *H*_*GMM*_, *H*_*RA*_ are used. If one treats a time series with GMM and the latter returns a zero value for *H*_*GMM*_, it means that the time series is stationary and its analysis should be made either by rescaled ranged analysis (RA), or any other method proper for analysis of stationary time series. Instead, if a time series is analyzed by RA and the latter returns Hurts exponent higher than one then it is not stationary and analysis should be made by GMM.

In the present work, viscoelastic and noise analyses of the approaching-retracting AFM-NI responses of *Ulocladium Chartarum* spores suggest the presence of biomemory effect in the cells functionality during external forcing, in agreement with previous works. ^1,2^ We define the response force, for pyramidal tip, for both approach and retraction parts of a FDC, and we extend the analysis in order to obtain in a single run both the viscoelastic scaling exponent and the scaling exponent of the power spectral density (PSD) that underlines the type of the environmental noise. The latter can operate as starting input in advanced mathematical modeling, fractional calculus, where knowledge of the environment’s noisy properties is mandatory.

### Power law rheology and Force Distance Curves under Linear Ramp

The response force, *f*(*h*), consists of the recorded values of the deflection signal with *h*(*t*) being the indentation depth. Assuming that a rigid indenter goes against to and/or penetrates a linear viscoelastic sample, *f*(*t*) and *h*(*t*) are related through convolution integrals, first introduced for spherical indenters, ^61^

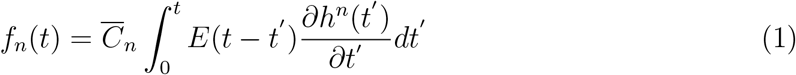

and

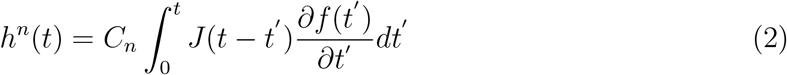

where 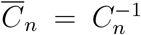, and the index *n* stands for the type of the indenter, with *n* = 1 for flat-ended cylindrical indenter with radius *R*, 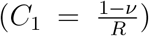, ^62^ *n* = 3*/*2 for spherical indenter,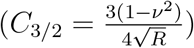, ^18^ and *n* = 2 either for conical indenter, 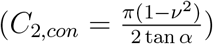, 63 or for four-sided pyramidal indenter with 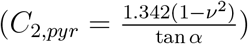. ^64^ The Poisson’s ratio is represented by *ν* and for incompressible materials takes the value of 0.5, and *α* is the average contact angle. *E*(*t*) and *J*(*t*) are the time dependent relaxation and creep function, respectively. By taking the in time-domain Laplace transform, 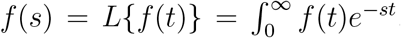, of each one of equations (1) and (2) and by setting *h*(0) = *f*(0) = 0, one can easily see that *E*(*s*)*J* (*s*) = 1*/s*^2^. Creep and relaxation function for viscoelastic materials follow a power-law behavior, that is, 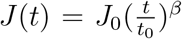, and 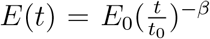. *E*_0_ and *J*_0_ satisfy the relation 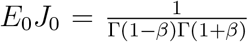, with 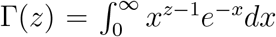 being the gamma function, notice that *E*_0_ is expressed in units of pressure (*N/m*^2^). Equations (1) and (2) hold when the contact area is an increasing function of time.^61^

#### Approaching phase

The indentation depth for constant velocity, *v*_0_, is a linear function of time and reads *h*(*t*)=*v*_0_*t*, for 0 *< t* ≤ *t*_*m*_ (loading or approaching phase), and *h*(*t*) = *v*_0_(2*t*_*m*_ − *t*), for *t*_*m*_ *< t* ≤ 2*t*_*m*_ (unloading or retracting phase). *t*_*m*_ is the time needed for the tip to reach its maximum penetration depth, where the phase changes from approach to retract. At this point, the direction of the velocity changes, but its speed is kept constant. For a typical AFM FDC of 1024 sampling points, *t*_*m*_ corresponds to 512 sampling points, and is converted to time units when multiplied by the minimum lag time, which is defined by the resolution of the machine. For fixed number of data points, and for pretty much constant maximum penetration depth, the surface anaglyph can lead to re-adjusting the initial position of the piezo, see Figure 1.

**Figure 1:**
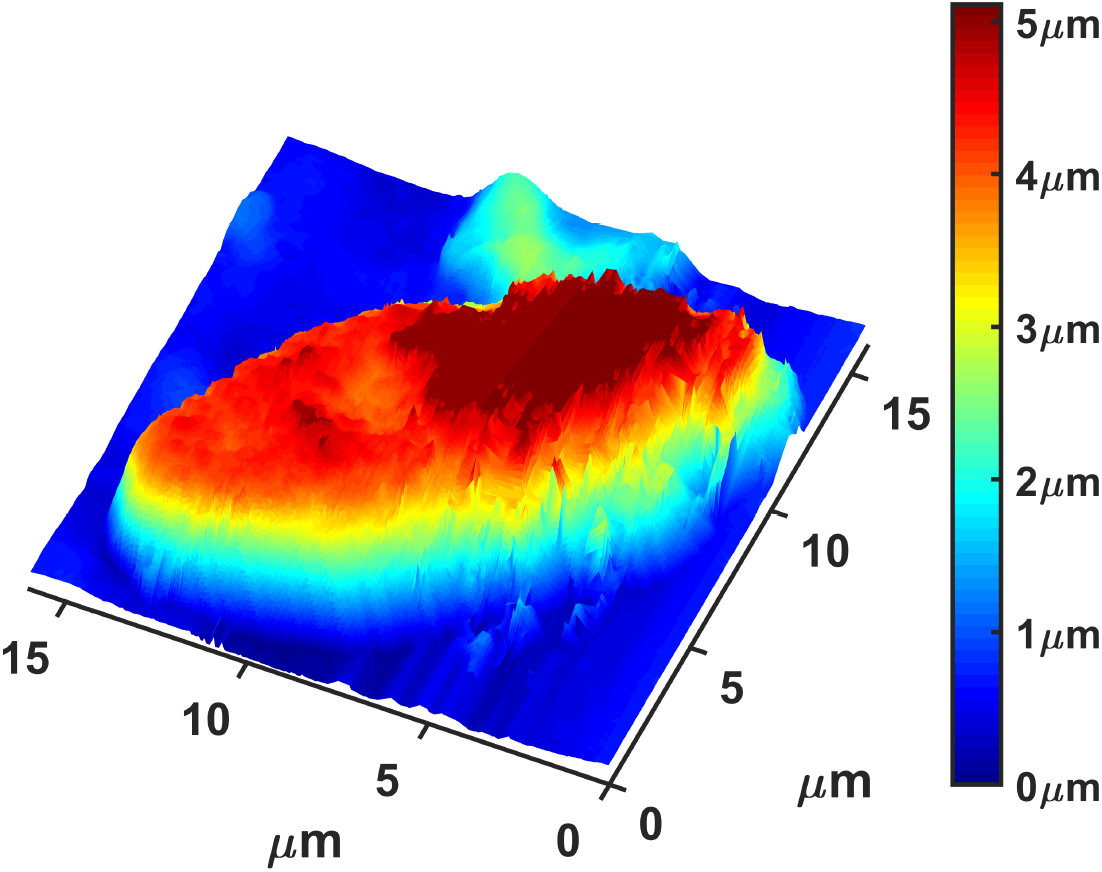
AFM image of an *Ulocladium Chartarum* single spore recorded under standard environmental conditions. Its surface is characterized by an intense changing landscape against whose the tip is moving with constant velocity *v*_0_ (towards/away)

For *h*(*t*) = *v*_0_*t* and by using eq.(1), one ends up with the recorded force, which reads

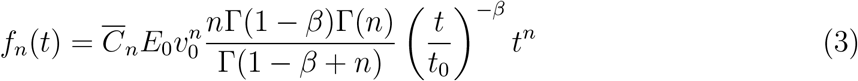

For a pyramidal type of indenter, *n* = 2, eq.(3) reads

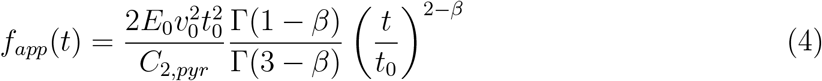

where *C*_2,*pyr*_ = 3.2, since we assume *α* = 17.5^*o*^ (the average contact angle), and *ν* = 0.5 (incompressible material).

#### Retracting phase

Eq.(4) holds for 0 *< t* ≤ *t*_*m*_ (approach) where the contact area monotonically increases as function of time.^61^ Eq.(4) can also be used for retraction, when the contact area decreases, under proper modification of time by following Ting’s method.^65,66^ The method assumes that there exist a time moment *t*^∗^, *t*^∗^ ∈ (0, *t*_*m*_), such that the contact area in the approaching phase is the same of the contact area in the retracting phase, *t* ∈ (*t*_*m*_, 2*t*_*m*_). So, the force can be obtained by equation (1) where we replace the upper limit of the integral from *t* to *t*^∗^(*t*), and then the link between the two time moments can be established by solving the following integral equation ^65^

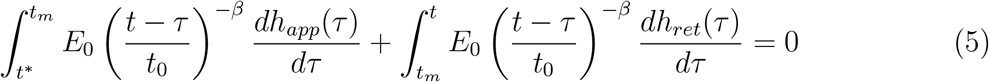

By replacing *h*_*app*_(*t*) = *v*_0_*t*, and *h*_*ret*_(*t*) = *v*_0_(2*t*_*m*_ − *t*) in Eq.(5) and by carrying out the integrals we find 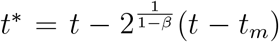.^21^ Notice that the value of *t*^∗^ depends only on the loading conditions and neither on the geometry of the tip nor on the thickness of the sample. The response force in the retracting phase is given by eq.(4) where we use *t*^∗^ instead of *t* and reads

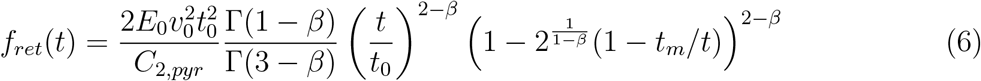

for *t* ∈ (*t*_*m*_, 2*t*_*m*_). The time derivative of eqs.(4) and (6) scales approximately as

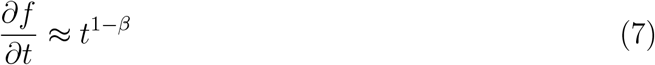

Corrections in eqs.(4) and (6) should be introduced to consider the influence of a solid support on the measured values of the response force. Such corrections are needed when cells, and in general samples that are measured, adhere on a surface (the solid support), usually glass whose Young’s modulus is orders of magnitude higher than that of cells. Back scattering effects originated from the solid support give a significant contribution to the force, especially when, the tip radius is comparable to the sample thickness. ^21,43,67^

### Treatment of Force Distance Curves as Time Series

FDC’s, experimentally recorded, can be considered as a sequence of values of either force, *F* (*n*), or distance, *h*(*n*), at hierarchically distributed time moments, *t*_*n*_, *n* = 1, 2, 3, .., *N* with *N* the maximum number of data points. Response forces are in the range of pN to *µN*, and are affected by environmental random forces that provide a stochastic contribution to the overall system. On the one hand, such random forces, contain information about the environment, and on the other hand, likely render questionable Young’s modulus values when they are the result of a direct fit of FDC’s to either eq.(4) or (6), vide infra.

### Generalized Moments Method (GMM)

A stochastic process can be memory-less, persistent, anti-persistent, where the characterization stands with respect to the kind of memory maintained by the process. If every new value of the stochastic sequence does not pose any dependence on its previous values then we call the process memory-less. Instead, if every new value depends on its previous values, then the process possesses memory. It is called persistent, when every new value likely follows the previous ones’ trend, and anti-persistent otherwise.

GMM is used to analyze non-stationary time series, ^56^ it is one of the most robust methods and works well even for short time series, ^50^ and it has successfully been applied in a several diverse fields.^51–55,68^ GMM uses the scaling of statistical moments of various orders including fractional ones. Briefly, GMM works as follows: It considers a time series of the form {*x*_*n*_} with *n* = 1, 2, …, *N*, where *N* is the total number of steps (measurements). If the minimum lag time is *τ* - the reciprocal of the sampling frequency - then the total length of the trajectory (time) is *T* = *Nτ*. If {*x*_*n*_} is a self-similar process then we expect that the time series, when zoomed in or zoomed out, will reveal the same patterns scaled by a certain amount, 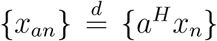, where 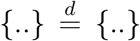 stands for the equality of finite dimensional distributions, and *H* ∈ (0, 1) is the scaling exponent also known as Hurst exponent. ^57^ We take the norm of the difference of {*x*_*n*_} at two distinct time moments, *t* and *s* with (*t > s*), and we write ||*x*_*t*_ − *x*_*s*_ || = ||*x*_1_ || || *t*^*H*^ − *s*^*H*^ ||. Furthermore, we consider three points *x*_*t*_, *x*_*s*_ and *x*_*t*−*s*_, their norms satisfy the inequality || *x*_*t*_ − *x*_*s*_|| ≤ || *x*_*t*−*s*_|| + || *x*_*t*_ − *x*_*s*_ − *x*_*t*−*s*_|| and by using || *x*_*t*_|| = || *x*_1_ || || *t*^*H*^ ||, we then end up with || *t*^*H*^ − *s*^*H*^ || ≤ || *t* − *s*|| *H* + || *t*^*H*^ − *s*^*H*^ − (*t* − *s*)^*H*^ ||, where the second term of the inequality goes as *s*^*H*^ for *s << t*. In this limit one can write, || *x*(*t*)−*x*(*s*) || ∼ | *t*−*s*|*H*, which for various moments of order, *q*, reads || *x*(*t*)−*x*(*s*) ||^*q*^ ∼ | *t*−*s* |^*qH*^. The latter has been extended to include the dependence of the Hurst exponent on the order of the moment, *H*(*q*) instead of *H*,^34,46,69,70^ since the various moments may not scale precisely by the same factor. The new exponent *z*(*q*) = *qH*(*q*) is called structure function. For discrete data sets the time difference *t* − *s* corresponds to a sliding window of length Δ, which must be small with respect to the total length of the time series.

#### First step

we construct time series characterized by different lag times, Δ, which contain the absolute change of the values between two points of the initial time series, let’s say *x*(*n*), that are apart by Δ:

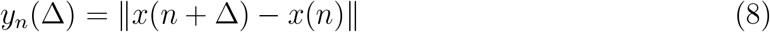

for *n* = 1, 2, …, (*T* − Δ)*/τ* and for 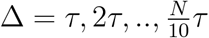. In order to have statistically reliable results, we define the maximum lag time as one-tenth of the maximum length of the original time series, *τ*_*max*_ = *N/*10, creating thus *N/*10 new time series of length (*T* − Δ) each.

#### Second step

We estimate the statistical moments of *y*_*n*_(Δ) according to

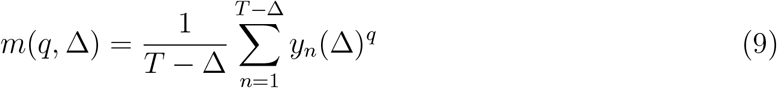

where also fractional values of the moment, *q*, are taken into account. We use only positive values of the moments.^71^ Moments in the range 0 *< q* ≤ 2 are responsible for the core of the probability density function (pdf), while moments higher than 2, *q >* 2, contribute to the tails of the pdf.^72^

#### Third step

we expect that the moments scale according to the elapsed time, Δ, as a power law

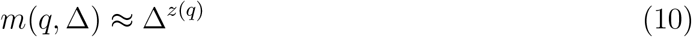

where *z*(*q*) is the structure function whose shape gives information on the stochastic mechanism(s) governing the motion. If the structure function is linear with respect to the order of the moment

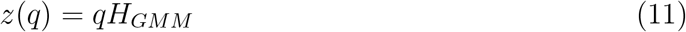

then, the process is mono-fractal, while if the structure function has a convex shape then the process is multifractal, see for details.^55,56^ Note that in eq.(11), *H* has been replaced by *H*_*GMM*_ to distinguish the analysis method.

### Rescaled Analysis (RA)

Let’s assume that *f*_*i*_ is a stationary time series, with *i* = 1, 2, 3, …, *N*. We divide the time series into *L* non-overlapping windows (sub-periods) of length Δ, 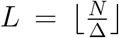. Δ provides the number of data points in a given sub-period that should be small with respect to *N*, and takes on the role of time when multiplied by the time lag. We fix its maximum value to 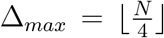, while its minimum value is set to Δ_*min*_ = 10. The number of the non-overlapping windows lies in the range 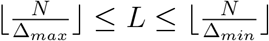. For each one of these windows we estimate the mean, 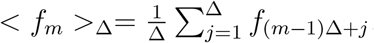, where *m* = 1, 2, .., *L*, and the standard deviation 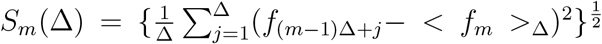. We create the profile 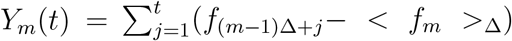. We estimate the distance *R*_*m*_(Δ)= *max*_1≤*t*≤Δ_{*Y*_*m*_(*t*)}-*min*_1≤*t*≤Δ_{*Y*_*m*_(*t*)}. We average all over the *L*-windows, 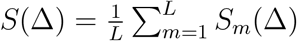, and 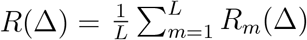, and we define the rescaled range (*R/S*)(Δ) which scales as^73^

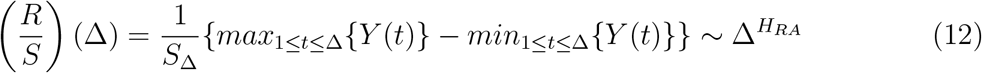

The quantity (*R/S*)(Δ) returns the rescaled distance between the maximum and minimum value of the time series of a given window of length Δ. According to the scaling described by eq.(12) this quantity is a monotonically increasing function of the length of the window. If in this description, *f*_*i*_ represents the differentiation of the response force recorded in AFM-NI then the scaling described by eq.(12) is the discrete analogue of the scaling described by eq.(7). This is true because of the time derivative of the response force (approach or retract), eq.(7), is a monotonically increasing function of time for 0 *< β <* 1.

Linear regression of equation (12) provides the exponent *H*_*RA*_

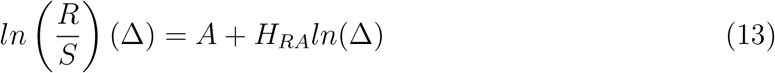

By equating the exponent of eq.(7) with the scaling exponent predicted by RA, eq.(13), we end up with

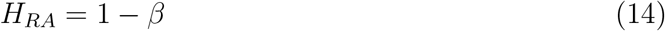

Equation (14) allows estimating the scaling exponent *β* whose value classifies the cell as elastic or liquid or viscoelastic.

For the construction of the time series *f*_*i*_ from the recorded FDC’s data we work as follows. In the approaching phase, the values 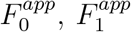 correspond to the response signal to the left of and at the contact point (CP), and the value 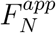 corresponds to the response force at *t* = *t*_*m*_. In the retracting phase, 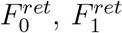 correspond to the values of the response force at *t* = *t*_*m*_ and *t* = *t*_*m*_ + *τ*, respectively, and 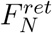 gives the value at the CP. A two-step pre-processing is required for further analysis; first we define the time series 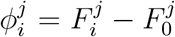 *i* = 0, 1, 2, .., *N* which describes the raw data shifted by the initial value, 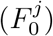, and second, we differentiate the sequences with respect to time, 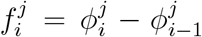, with *i* = 1, 2, .., *N*.

The new time series, 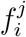, is the derivative of the response force multiplied by *τ*. On the other hand, the accumulation of 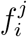 up to a certain time moment gives the shifted response signal, 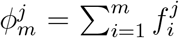. For 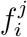 being stationary, RA is used and the viscoelastic exponent is provided by eq.(14). For non-stationary 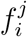, GMM is used and the viscoelastic exponent is again provided by eq.(14) where we replace *H*_*RA*_ with *H*_*GMM*_.

Parallel to environmental noise an instrumental AFM noise, which is a function of instrumental, thermal, acoustic, electronic and quantum noises e.t.c., is also present. These additional noise contributions are characterized by different time scales, and probably die out at the time scale defined by the resolution of the instrument. Constant contributions arising from an AFM noise cannot affect our analysis because of the proposed methodology is based on differences between values of subsequent steps that cancel out constant contributions. If, however, these components of AFM noise are not constant in time and turn the measured signals (their time derivatives) into multiplicative ones (i.e. non-stationary), then their nature can be identified by the GMM, which returns structure function of convex shape, vide infra.

### Power Spectral Density (PSD)

A widely accepted measure for the classification of stationary time series is its PSD, while for non-stationary time series this measure is questionable. ^74^ In many phenomena, PSD scales as

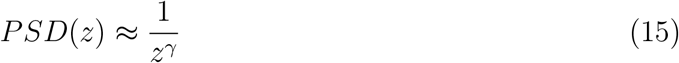

where *z* is the variable in the frequency domain, and *γ* the scaling exponent whose value defines the color of a stochastic sequence. Colors are well-defined for −1 ≤ *γ* ≤ 1, white for *γ* = 0, blue for *γ* = −1, pink for *γ* = 1. For *γ <* −1 or *γ >* 1, colors exist, for example, purple for *γ* = −2, red or brown for *γ* = 2 and black for *γ >* 3. Criteria for color classification are not tight, so a signal with −1.3 ≤ *γ* ≤ −0.5 can be characterized as bluish, in the range 0.5 ≤ *γ* ≤ 1.5 as pink or flicker. For time series defined in the time domain the scaling of their PSD do not exceed the value of 2.^74^ Eq.(15) is an approximation good only for low frequencies. In practice, eq.(15) provides adequate scaling exponents for real-life time series, only if the scaling holds true for at least two decades in the frequency domain. ^60^ Spectral methods accurately predict scaling of synthetic time series produced in the frequency domain, while for those produced in the time domain, half of the spectra estimates deviate significantly for the nominal value of *γ*.^75^ The sign of the exponent characterises a process as anti-persistent, negative (purple, blue), and as persistent, positive (pink, red, black). Additionally classification is made regarding stationary or non-stationary nature of the process: fractional Gaussian noises, fGn, (stationary processes) for −1 *< γ <* 1, and fractional Brownian motion, fBm, for 1 *< γ <* 3 (non-stationary process). Notice that fBm is the integration of fGn up to time *t*. Assuming either fGn, or fBm as the kind of the underlying stochastic process then there is a direct connection between the Hurst exponent and the power spectrum scaling exponent^60^

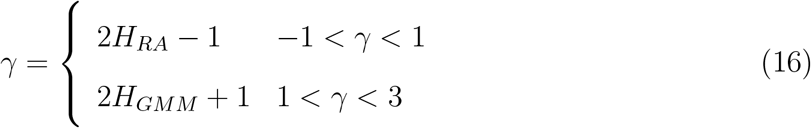

In equation (16) the subscript refers to the analysis method used to obtain the Hurst exponent, RA for stationary noises and GMM for non-stationary. By using the path of analysis described above one can easily find the scaling exponent describing the viscoelastic properties of the cell, eq.(14), as well as the scaling exponent providing information about the noise properties of the cell, eq.(16), assuming the existence of fractional Gaussian noises.

## Materials and Experimental Part

We test our model on rubber that is a “non-living” material; Polydimethylsiloxane (PDMS) has been used as sample. Two sets of four AFM-NI experiments each have been carried out with the same set up of the experiments carried out in spores, and two different control conditions. The first one, considers velocity of penetration, *u* = 0.029*µm/sec*, very similar to velocity of penetration in spores, and the second one considers much higher velocity of penetration, *u* = 0.49*µm/sec*, see below. A detailed analysis of these experiments is given at Section I of the Supporting Information (SI), see also Figures S1a, S1b, S2, S3a, and S3b as well results listed in Table S1. The time derivatives of the measured deflection signals (Fig.S1a and Fig.S1b) correspond to non-stationary time series and analysis has been made by GMM. For the first control condition, GMM delivers viscoelastic exponents in the range [0.758 – 0.850], for approach, and in the range [0.734 - 0.758], for retraction. Both phases describe a liquid-like behavior of the “non-living” material. For the second control condition the corresponding values are [0.362 - 0.415], for approach, and [0.316 - 0.342] for retraction, and the “non-living” material behaves as a viscoaelastic one. We found by using two control conditions of substantial difference in the velocity of penetration that the higher the velocity of penetration the more elastic the material appears. ^21^ We also used eqs (4) and (6) to fit directly the measured deflection signals for all experiments. Eq.(4) fits well the approach phase for both control conditions returning exponents in very good agreement with what has been obtained by GMM, see Table S1. On the contrary, fittings with eq.(6) (retraction) return values of *β* reduced by at least a factor of 4(2) for 0.029(0.49) */mum/sec* with respect to the approaching phase and to what is obtained by GMM. This inconsistency may due to additional contributions e.g. drift and/or feedback electronics, which challenge the assumption of a monotonic decreasing contact area, a necessary condition for application of eq.(6). These contributions can not affect the GMM since the latter uses the absolute changes between two observables.

Living *Ulocladium Chartarum* were cultivated on potato dextrose agar (Merck, *pH* = 5.6) at 298 K. Part of the culture was uniformly spread over an area of 250 *mm*^2^ on the coverslip substrate under an optical metallographic microscope, (Leica DMRX). The spore cells were left to dry on air after removing traces of humidity with a paper filter. Caution was taken to form monolayers of spores preventing inner spore shielding. The nanoindentation was carried out with the same type of cantilever, at ambient conditions using phosphorus-(n)-doped silicon cantilever (Bruker RTESPA-300) having a nominal spring constant and resonance frequency of 40 *N/m* and 300 *kHz*, respectively. A stiff cantilever has been chosen to ensure that the tip penetrates inside the hard *Ulocladium Chartarum* membrane.^76^ This choice may reduce the hysteresis between approach and retraction, see Figure 2.

**Figure 2:**
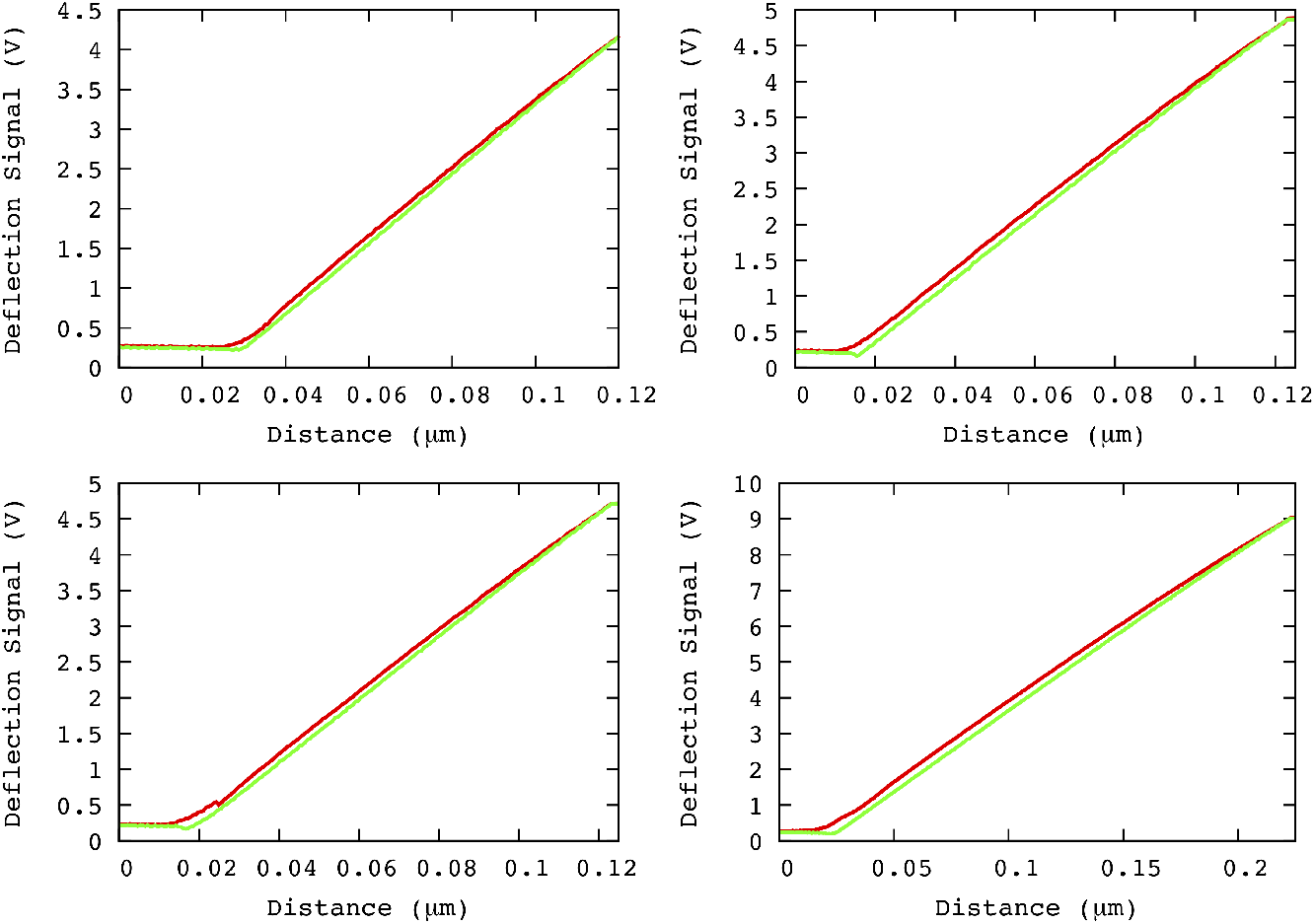
Color code: red for approach and green for retraction. A part of Force Distance Curves for some typical experiments analyzed in this study are depicted.

FDCs were recorded at different points, regularly distributed over pre-selected areas in the center of the cell, and away from the spore edges avoiding thus artefacts introduced by cell boundaries. We have conducted thirteen experiments, and three constant velocities namely, 0.0248/0.0289/0.0331 *µm/sec*, have been used. The maximum penetration depth lies in the range of 100 - 133 *nm*, but there is also a value of 83 *nm* as well as of 207 *nm*.

Fig.2 shows some of FDC’s in AFM-NI experiments, note that in all experiments we used only one type of cantilever. The deflection signal in Volts can be turn to force by multiplying the voltage values by a proper conversion factor, which may also present dependency on the geometry of the laser beam. The treatment is independent on voltages or forces since the beta value involves differences and does not dependent on the conversion factor. The recorded curves consist of 1024 sampling points, and thus the phase changes from approach to retraction at the time moment *t*_*m*_, which corresponds to 512 sampling point. The curves are converted to time units when multiplied by the minimum lag time (reciprocal of the sampling rate), which is approximately equal to 23.6 *ms*.

## Results and Discussion

The approach and retraction pathways do not coincide, Fig. 2. Approach and retraction evolve under constant velocity, which has been kept slow with respect to the time-scale of molecular re-organisation. The two processes, therefore, form a pair of dynamical processes that do not coincide and evolve near equilibrium. In principle, the response mechanisms governing these processes in living cells can differ and, accordingly, differentiate the response of a biological system under the stimulus of the same mechanical object. Approach and retraction are treated separately for each experiment. To estimate the scaling exponent *β*, we use the curves depicted in Fig.3a/Fig.3c, which are parts of the overall FDCs and correspond to measurements taken with the tip in contact with the sample. These parts of a FDC form discrete time series of equidistant points, 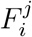, *i* = 0, 1, 2, .., *K*, where every pair of two consecutive points is separated by the minimum time lag, *τ*, and *K* provides the maximum number of data points of which the analysed part of the curve is consisted of. The obtained time series are non-stationary, see Figs. 3a and 3c.The index *j* stands for the phase and takes two values, *j* = *app* or *j* = *ret* for approach and retraction, respectively.

**Figure 3:**
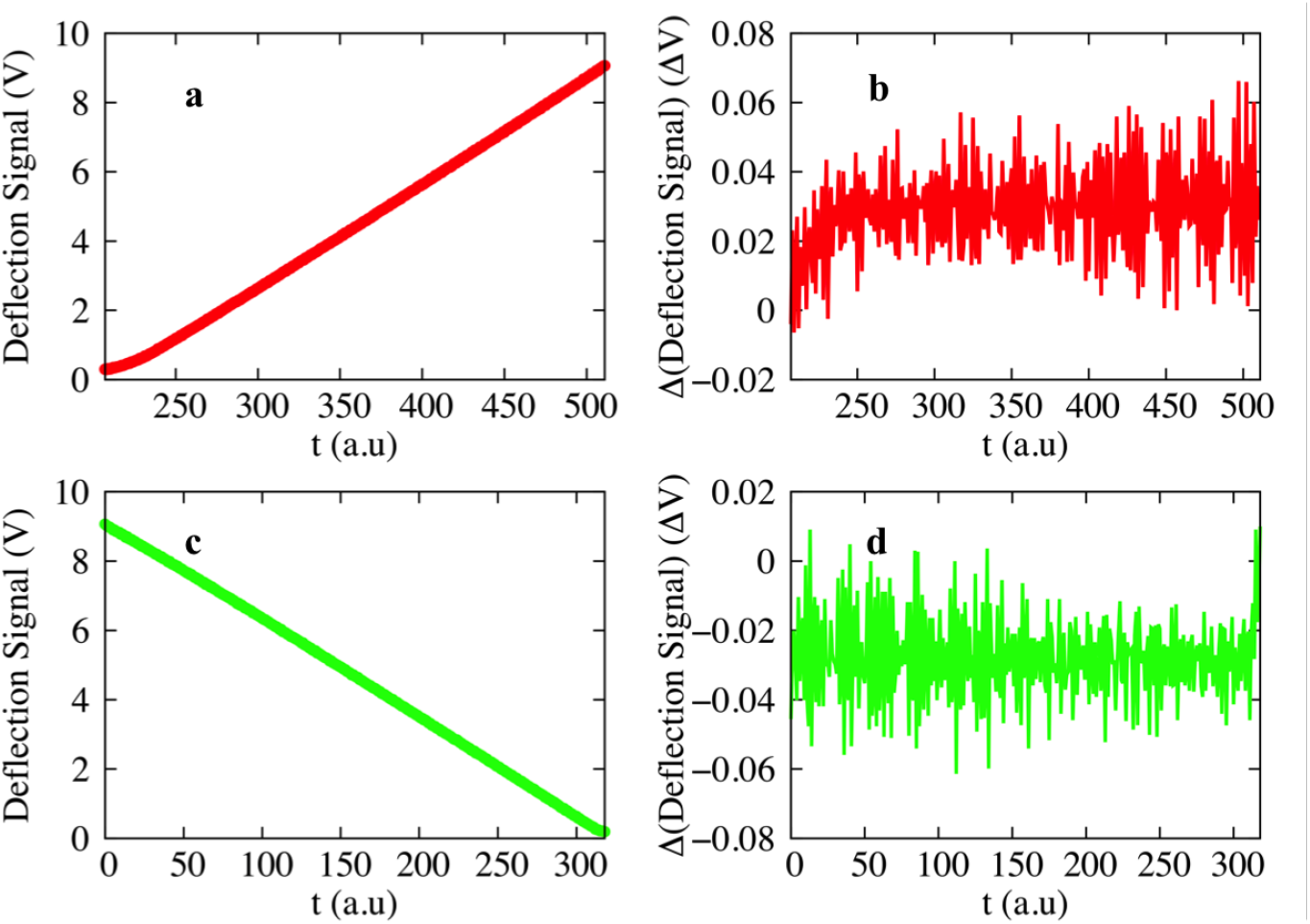
Color code: approach (red) and retraction (green). Recorded deflection signal in a typical AFM-NI experiment is illustrated in a) and c) where data after/before the contact point for approach/retraction are used. The differentiation of the recorded signal is given in b) and d) for approach and retraction respectively. These data sets are studied by using GMM and RA.

We apply GMM in order to verify if 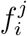 is stationary or not,Figs.3b and 3d. For all experiments, for both approach and retraction, GMM analysis returns a zero value for *H*_*GMM*_.

It implies that 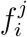 are stationary and their further analysis is made using RA by means of equation (12). Linear regression of eq.(12) returns the values of *H*_*RA*_, eq.(13). Figure 4 shows the best fits as well as the estimated exponents for four experiments. The obtained Hurst exponents, *H*_*RA*_, differentiate pathways for approach and retraction. For the approaching phase, and for similar maximum penetration depths of about ∼ 100 *nm* (only in one experiment the maximum penetration depth is ∼ 200 *nm*), we obtain values of *H*_*RA*_ lying in the range 0.48 ≤ *H*_*RA*_ ≤ 0.75. For the same experiments in the retracting phase the returned value of *H*_*RA*_ lies in the range 0.15 ≤ *H*_*RA*_ ≤ 0.33, see Table 1.

**Table 1:** Approach/Retraction process for different points regularly distributed over pre-selected areas in the center of a *Ulocladium Chartarum* spore. The 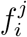 time series created by the raw data are first treated by using GMM. The method delivers *H*_*GMM*_ = 0.0 for all experiments and for both phases proving thus the stationary nature of the 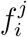. The same time series are then treating by RA, which delivers the scaling exponents, *H*_*RA*_, of the derivative of the response force by means of equation (14), the standard error of estimate is also provided. By using eqs. (14) and (16) we obtain the scaling exponents characterizing the viscoelastic material and the PSD, *β* and *γ*, respectively.

| Exp. | $u$ ($\mu m/sec$) | Approach | | | Retraction | | |
| --- | --- | --- | --- | --- | --- | --- | --- |
| | | $H_{RA}$ | $\beta$ | $\gamma$ | $H_{RA}$ | $\beta$ | $\gamma$ |
| 1,2 | 0.0248 | $0.48 \pm 0.025, 0.61 \pm 0.014$ | 0.52, 0.39 | -0.04, 0.22 | $0.19 \pm 0.031, 0.23 \pm 0.015$ | 0.81, 0.77 | -0.62, -0.44 |
| 3,4 | 0.0248 | $0.55 \pm 0.015, 0.60 \pm 0.016$ | 0.45, 0.40 | 0.10, 0.20 | $0.25 \pm 0.018, 0.15 \pm 0.016$ | 0.75, 0.85 | -0.50, -0.70 |
| 5,6 | 0.0289 | $0.66 \pm 0.024, 0.63 \pm 0.022$ | 0.34, 0.37 | 0.32, 0.26 | $0.19 \pm 0.013, 0.24 \pm 0.024$ | 0.81, 0.76 | -0.62, -0.52 |
| 7,8 | 0.0289 | $0.62 \pm 0.027, 0.52 \pm 0.021$ | 0.38, 0.48 | 0.24, 0.04 | $0.27 \pm 0.035, 0.20 \pm 0.024$ | 0.73, 0.80 | -0.46, -0.60 |
| 9,10 | 0.0289 | $0.61 \pm 0.030, 0.52 \pm 0.020$ | 0.39, 0.48 | 0.22, 0.04 | $0.31 \pm 0.026, 0.24 \pm 0.026$ | 0.69, 0.76 | -0.38, -0.52 |
| 11,12 | 0.0289 | $0.59 \pm 0.027, 0.58 \pm 0.035$ | 0.41, 0.42 | 0.18, 0.16 | $0.24 \pm 0.031, 0.33 \pm 0.017$ | 0.76, 0.67 | -0.52, -0.34 |
| 13 | 0.0331 | $0.75 \pm 0.030$ | 0.25 | 0.50 | $0.23 \pm 0.023$ | 0.77 | -0.54 |

**Figure 4:**
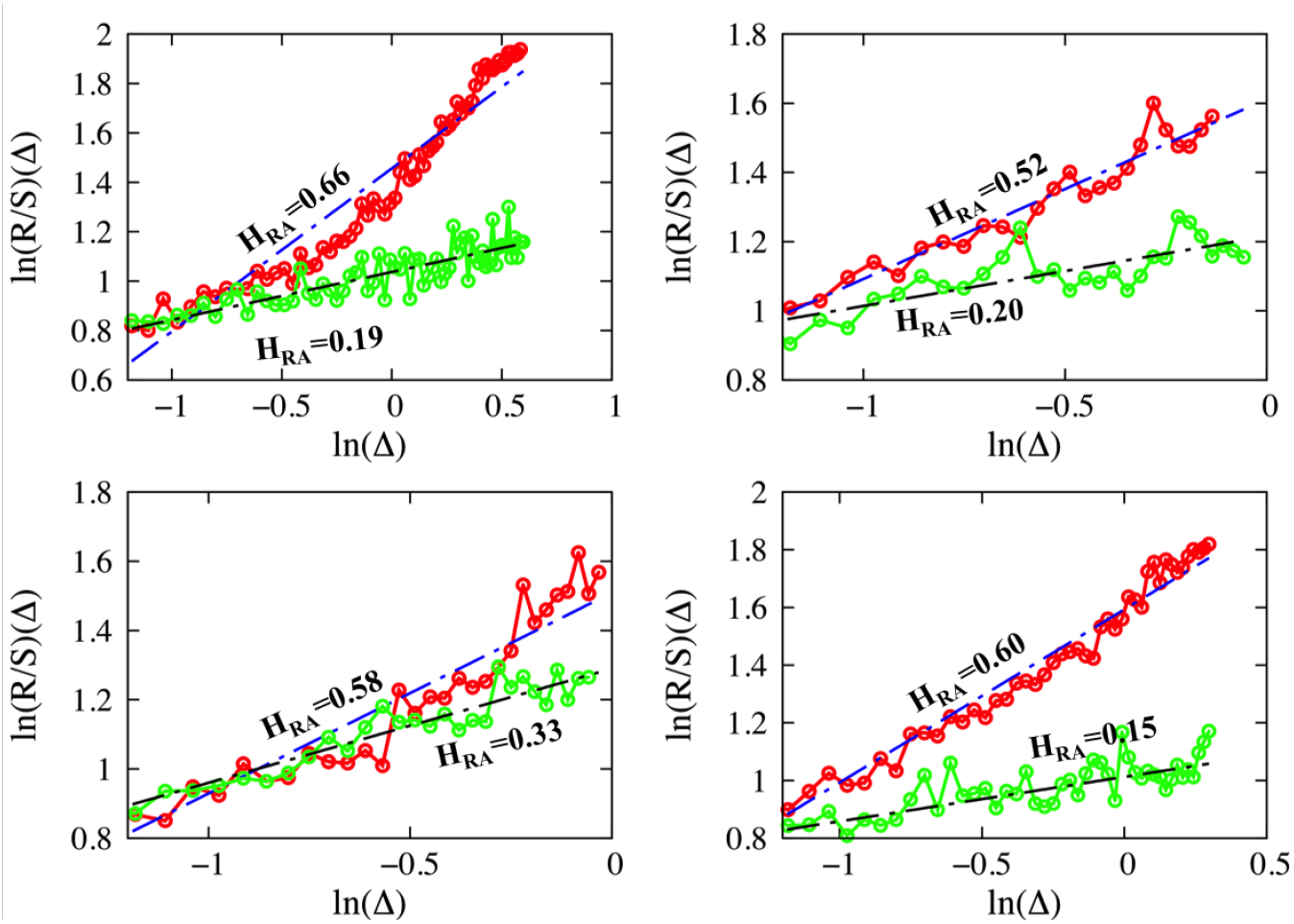
Four different experiments are illustrated. Each panel describes the application of RA for the approach and the retraction phase of the same experiments. Color code: red for approach and green for retraction. Fittings have been performed by using equation (13), dashed blue/black lines for approach and retraction, respectively. The slope of each fit returns the Hurst exponent, *H*_*RA*_, which is also depicted in the graphs.

The scaling exponents characterize the viscoelastic properties of the cell, are obtained by using eq.(14), their values are listed in Table 1, and illustrated in Fig.5. In the approaching phase, the Hurst exponents are greater than or equal to 0.48, which is typical either of uncorrelated processes for values close to 0.5 or of slightly persistent processes for larger values. It means that the cell, in most of the cases, attempts to counterbalance nearly perfect the effect of the tip. Additionally, the conjugated scaling exponents, *β*, eq. (14), lie in the range of 0.25 ≥ *β* ≥ 0.52. The values align with the literature where *β* is often found to lie in the range of 0.1-0.3. ^27,28^ Values of *β* close to 0.4 have been reported for fibroblast (NIH 3T3),^21^ and values close to 0.5 have been found for maximum penetration depth deeper than 1 *µm* ^31^ (not reach in the current study). On the other hand, increased values of *β* and close to 0.5 have been associated with regions of the cell in the periphery of the centre. The values of *β* are somewhat scattered, and they do not depend on the values of the low velocities used here. ^21^ Exception is the single value for the control condition 0.033 *µm/sec*, which might introduce a dependency condition, and its statistical significance possibly remains to be clarified in future experiments. A comprehensive microscopic interpretation of what the scaling exponent *β* represents is still missing, albeit that it has been proposed that *β* represents the turn-over dynamics of cytoskeletal proteins and cross-linkers, including myosin motor activity. ^27^ Cytoskeletal protein dynamics is essential for contraction and locomotion, ^77^ and has been reported to be higher in peripheral areas of the cell such as in the lamella and the lamellipodium, resulting in increased values of *β*. The scattered values of *β*, Fig.5, for similar penetration depths likely indicate cell inhomogeneity. It has been reported that there is no evidence for a dependence of the value of *β* on the depth of penetration, ^31^ an argument also satisfied in this study for the experiment number 5, where the penetration depth is almost the double with respect to the other ones.

**Figure 5:**
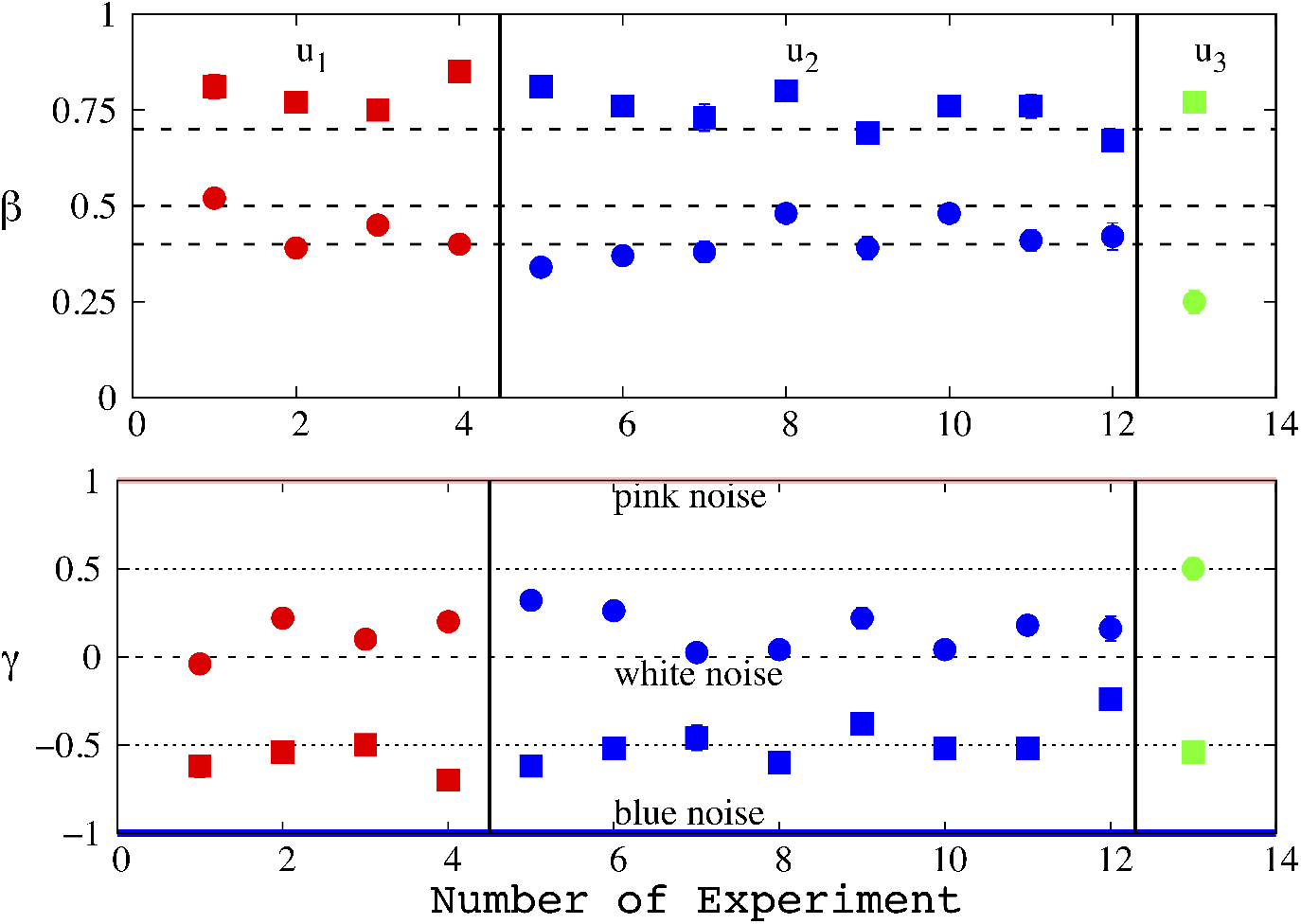
Color code to assist the eye for different constant loads: red for *u*_1_=0.0248 *µm/sec*, blue for *u*_2_=0.0289 *µm/sec*, and green for *u*_3_=0.0331 *µm/sec*. Symbols: circles for the approaching phase and squares for the retracting one. Top: scaling exponents, *β*, of the viscoelastic properties of cell. Bottom: Scaling exponent, *γ*, of the power spectrum. Three horizontal lines stand for some well-defined colors; blue for *γ* = −1, white for *γ* = 0, and pink for *γ* = −1.

The retracting phase of each experiment reveals an entirely different picture. The values of the Hurst exponent lie in the range of 0.15 ≥ *H*_*RA*_ ≥ 0.33, suggesting an anti-persistent process, where each step likely goes in the opposite direction of the previous one. It means that the tip experiences forces that oppose the predefined motion imposed by the tip and are of the same order of the latter. These forces are likely to be of capillary nature, caused by liquid filling of the path created by the tip during the penetration. The liquid nature of the path is corroborated by the values of the scaling exponent *β* lying in the range 0.67≥ *β* ≥ 0.85, which is typical of liquid-like materials. ^27,28,31^ Different responses of approaching retracting AFM-NI curves are associated with biomemory effects of cells for tracing maximum viability lines. External stress is responsible for exuding intracellular substances on the cell wall and a change in the intracellular environment for protecting the cell.^1,2,78–80^

For stationary processes the value of *γ* underlines the color of the process. *γ* can be obtained either by using eq.(15) or by eq.(16). Application of eq.(15), which requires a direct transformation of the differentiated data to Fourier space and then fitting with eq.(15), presents significant standard error of estimate, see Section II of Supporting Information (SI). We obtained the scaling exponents by using eq.(16). In the approaching phase, we find 0*< γ* ≤ 0.5, persistent noises and accordingly during this phase the cell retains a memory. On the other hand, the retracting phase is described by −0.7≥ *γ* ≥-0.34, bluish type of noises, values describe anti-persistence process again in line with properties underlined by the values of *β*. Bluish noises have been reported in literature as noises patterns used by retina cells to yield visual resolution.^81^ For both, approach/retraction the time derivative of the response force is described as fractional Gaussian noise, obtained values of *γ*, see Fig.5. This finding is introduced here for the very first time and could be exploited in the modelling of the AFM-NI motion, for instance, in terms of a fractional Langevin type equation where the proper form of the environmental noise must be chosen. Such a strict mathematical description of the cell response under an AFM-indenter of pyramidal shape can provide further information on the microscopic nature of the scaling exponent *β*. We leave this task for a future work.

## Conclusion

In summary, a robust methodology for the analysis of AFM-NI force distance curves is proposed, which can also be extended to incorporate contributions from the solid support. The analysis of experiments conducted in living *Ulocladium Chartarum* spores shows that approaching and retracting phases are truly different processes. Their different nature appears; a) by the scaling exponents describing their viscoelastic properties, and b) by the scaling exponents of their power spectral density, which is connected to the type of the environmental noise. In the approaching phase the cell presents a viscoelastic behaviour similar to what has already been reported in the literature. The process is persistent underlining a synergic action of the inner components of cell opposing to the motion of the tip. The retracting phase corresponds to an anti-persistent process and displays characteristics of a liquid-like material, which interacts with the tip by forces that are likely of capillary nature. These forces originate on the release of proteins and bio-substances, triggered by a mechanical stimulus, that fill the path created by the tip, and indicate a biomemory response of cells to local mechanical stress as the one imposed by the AFM tip during indentation. The environmental noise is bluish for the retracting phase and persistent, quasi flicker, for the approaching one.

## Supporting information

SI in the main text

## Supporting Information

Supporting Information material contains two sections. At the first one, analysis of the deflection signals, for two control conditions, measured by using “non-living” material is presented, and it is accompanied by figures S1a, S1b, S2, S3a and S3b as well as by the Table S1, where all findings are listed. The second section contains the Table S2, where the power spectrum scaling exponents, obtained either directly by linear regression of equation (15), or indirectly by using eq.(16), are listed.

## Acknowledgement

The work has been funded under the frame of the projects “ELI – LASERLAB Europe Synergy, HiPER and IPERION-CH.gr” (MIS 5002735) which is implemented under the Action “Reinforcement of the Research and Innovation Infrastructure”, funded by the Operational Programme “Competitiveness, Entrepreneurship and Innovation” (NSRF 2014-2020) and co-financed by Greece and the European Union (European Regional Development Fund and “Advanced Materials and Devices” (MIS 5002409) which is implemented under the “Action for the Strategic Development on the Research and Technological Sector”, funded by the Operational Programme “Competitiveness, Entrepreneurship and Innovation” (NSRF 2014- 2020) and co-financed by Greece and the European Union (European Regional Development Fund).

## Graphical TOC Entry

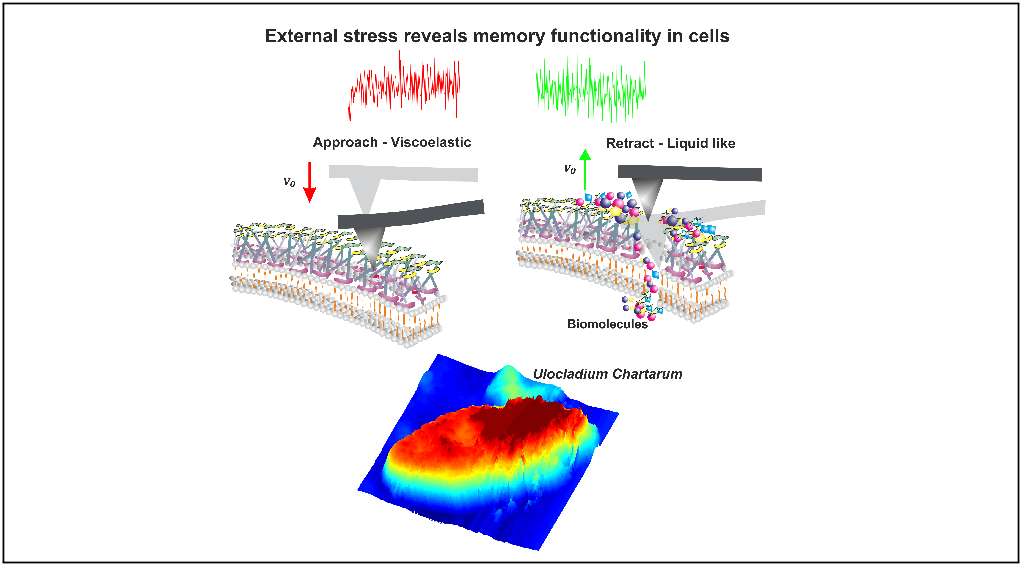

