## Supplementary material for "Viscoelasticity and Noise Properties Reveal the Formation of Biomemory in Cells": SI in the main text

**for**


### Section I.

In this section the viscoelastic exponent, β, of a rubber "non-living" material is obtained. We conducted two sets of four AFM-NI experiments at room temperature where Polydimethylsiloxane (PDMS) was used as a sample. In the first set of experiments approach and retraction rates are similar to that have been used in spores ~0.029 μm/sec, while in the second set of experiments, approach and retraction rates are equal to ~ 0.49 μm/sec (~17 times higher than the first control condition). PDMS is widely used in bio-transducers due to its biocompatibility and its mechanical compliance.[1] Viscoelasticity is an inherent property of PDMS, and it changes with loading rates and exposition times on loads.[2] For a PDMS sample, the viscoelastic exponent depends on the chain length and the concentration of the precursor material.[3] It has been also reported that for a scarcely cross-linked PDMS at room temperature the viscoelastic exponent presents a cross over point from low (β=0.3) to high (β=0.6) frequencies.[4]

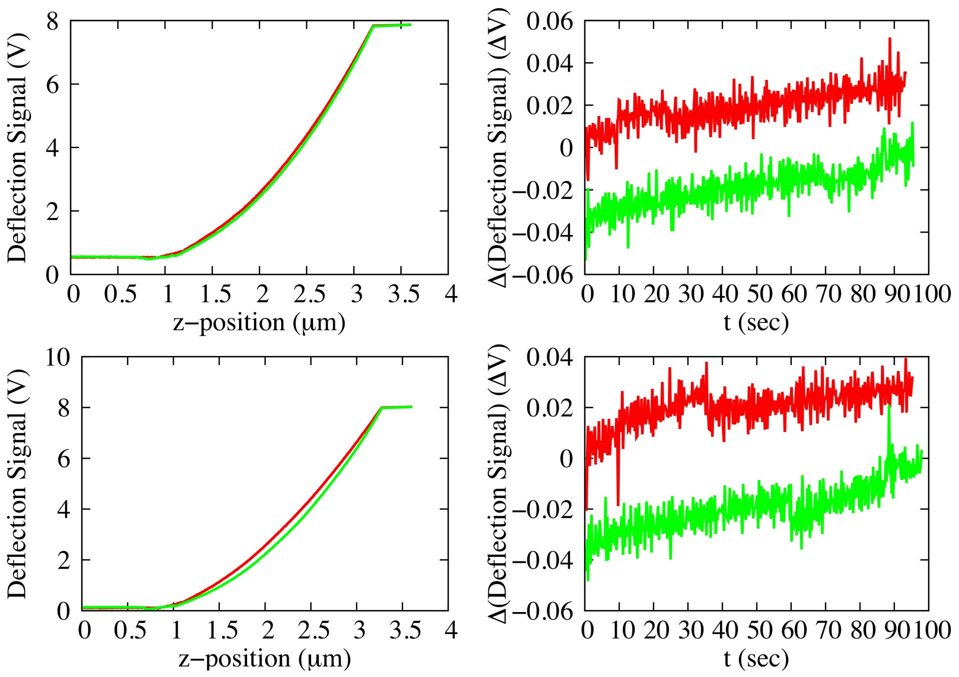


**Figure S1a. (Control condition, u=0.029 μm/sec)** Upper and lower panels: Left, Force Distance Curves (FDC) for two out of four experiments conducted in PDMS. Right, the differentiation of the recorded deflection signal with respect to time for the signals depicted on the left. For the differentiation, we considered only the points to the right/left of the contact point (CP) for approach/retraction respectively. Color code: red for approach and green for retraction.

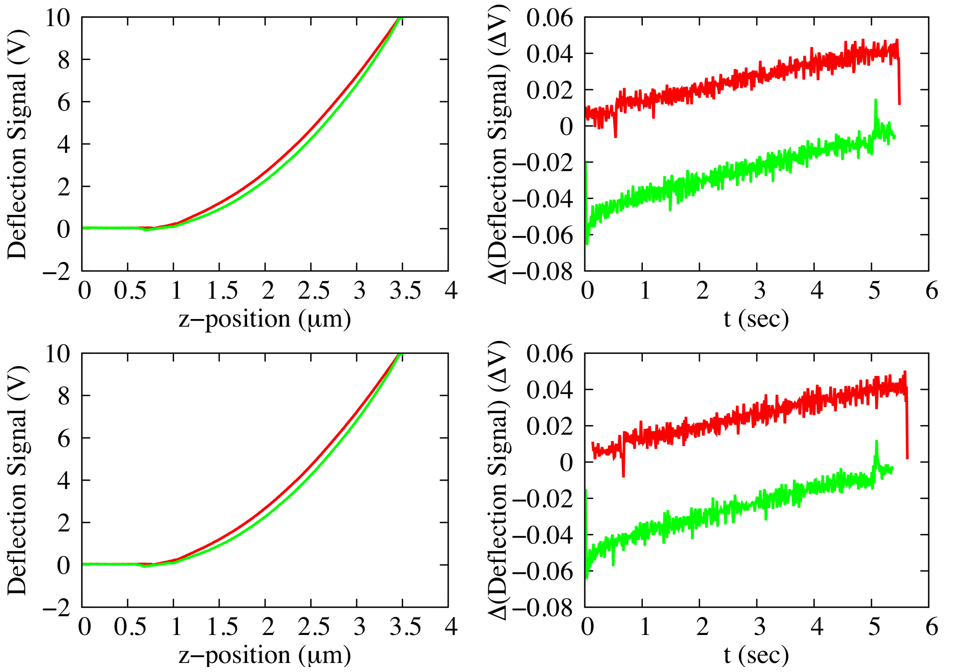


**Figure S1b. (Control condition, u=0.49 μm/sec)** Upper and lower panels: Left, Force Distance Curves (FDC) for two out of four experiments conducted in PDMS.

Right, the differentiation of the recorded deflection signal with respect to time for the signals depicted on the left. For the differentiation, we considered only the points to the right/left of the contact point (CP) for approach/retraction respectively. Color code: red for approach and green for retraction.

A direct estimate of the scaling exponent can be made by fitting eq.(4) of the main text to the raw data (see Figure S1a and S1b left panels) for the approach phase, and by fitting eq.(6) for the retraction. The experimental curves as well the corresponding fittings for each phase are depicted in Figure S2, and the obtained scaling exponents are listed in Table S1.

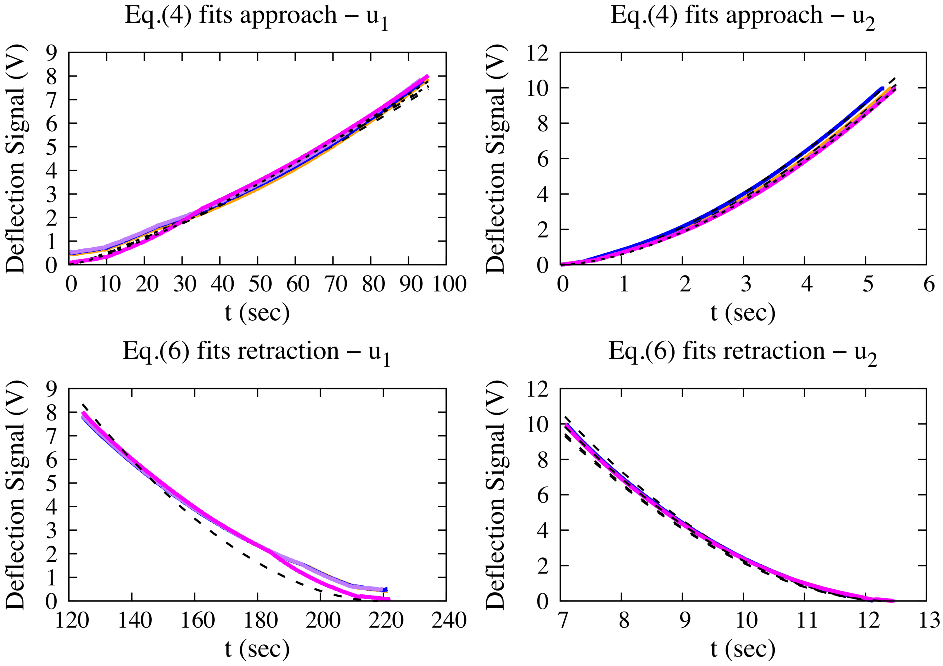


**Figure S2**. Recorded deflection signals in AFM-NI versus time; top: approach for two different control conditions ($u_l$=0.029 μm/sec and $u_2$=0.49 μm/sec), bottom: retraction for the same conditions. The black lines are the fittings.

The scaling exponent depends on the control condition, the higher the velocity the more elastic the material. In the approaching phase and for the first condition, $u_l$=0.029 μm/sec, the scaling takes values in the range [0.758 – 0.850] highlighting a liquid like material. For the same condition, in the retraction phase the scaling exponents take constantly values lower than 0.2 and close to 0.18 pointing to an elastic material. For the second condition, $u_2$=0.49 μm/sec, the scaling exponents take values in the range [0.331 – 0.416], while for retraction the scaling exponents have pretty much the same value for all experiments, (β ~0.16). The findings indicate that for a much higher velocity of penetration, the sample material behaves as a viscoelastic material, which, in the retraction phase, seems to have become more elastic because of the reduction of the scaling exponent by a factor of two.

We apply our method on the recorded FDC and compare the results with the scaling exponents returned by a direct fit of eqs (4) and (6). Fits with eq.(4) return values in very good agreement with what we have obtained by the proposed method in this article. GMM (or RA) uses as input the differentiated signals of the right panels of Figure S1. We demonstrate the way of selecting the correct analysis method for a given time series starting our analysis by using RA. The obtained Hurst exponents ($H_{RA}$) are listed in Table S1 (colored in red), and for each phase and for all experiments take values greater than 1. These values, and for every $H_{RA}\geq 1$, indicate that the underlying time series is non-stationary and the analysis method, RA, is not proper. GMM is a proper method for analysis of non-stationary time series and by following the steps described by eqs (8), (9), and (10) of the main text we obtain the structure functions whose forms are illustrated in Figures S3a and S3b.

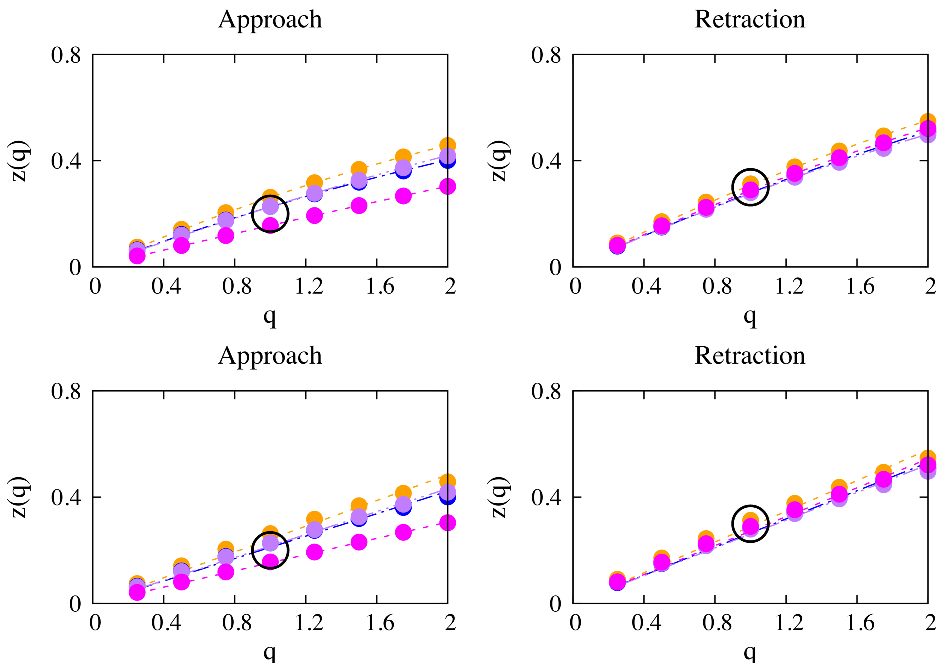


**Figure S3a**. (Control condition u=0.029 μm/sec) The structure function z(q) versus the order of the moment, q, for both approach and retraction. Upper panels: $z(q)=Hq-C(q^2-q)$, Lower Panels: $z(q)=Hq$ (eq.11 of the main text).

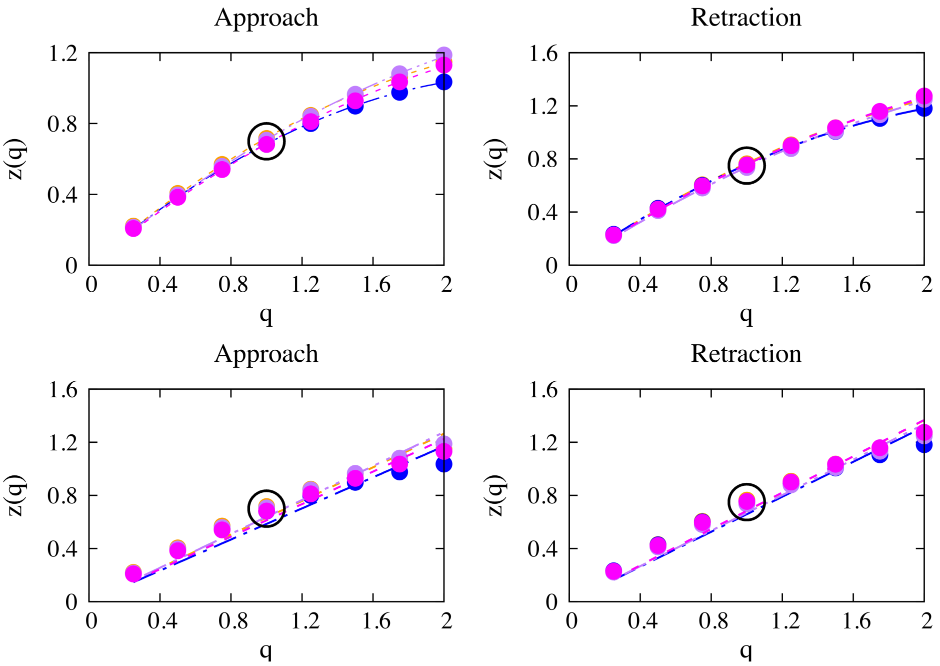


**Figure S3b**. (Control condition u=0.49 μm/sec) The structure function z(q) versus the order of the moment, q, for both approach and retraction. Upper panels: $z(q)=Hq-C(q^2-q)$, Lower Panels: $z(q)=Hq$ (eq.11 of the main text).

The structure functions for the two control conditions are illustrated in FigS3a (u=0.029 μm/sec), and FigS3b (u=0.49 μm/sec). The values of the structure function have been fitted by eq.(11) of the main text, lower panels, and by the form $z(q)=Hq-C(q^2-q)$ which describes the viscoelastic environment as including the presence of another mechanism acting multiplicatively.[5] By employing our hypothesis of the viscoelastic environment, eq.(11) returns Hurst exponents ($H_{GMM}$), listed in Table S1, whose connection with the scaling exponent is made through eq. (14), and the values of these exponents are listed in Table S1. In the approach phase, the agreement of the values of β's obtained by fitting eq.(4) and by $H_{GMM}$ is excellent, values in bold in Table S1. For the retraction phase, the β's obtained by GMM have constantly lower values with respect to corresponding values of the approach phase obtained by the same method.

**Table S1**: Viscoelastic exponents obtained by using eqs (4) and (6) of the main text, or by using eq.(14) of the main text where the value of the Hurst exponent is returned by the use either of GMM or of RA. The value of the Hurst exponent is listed for each experiment and for each phase.

| u=0.029 μm/sec | | | | | | |
|---|---|---|---|---|---|---|
| Exp | Approach<br>β | Retraction<br>β | Approach<br>β | Retraction<br>β | Approach | Retraction |
| | Eq.(4) | Eq.(6) | GMM (eq.14) | GMM(eq.14) | $H_{GMM}$ / $H_{RA}$ | $H_{GMM}$ / $H_{RA}$ |
| 1 | **0.742**±0.008 | <0.2 | **0.758**±0.005 | 0.734±0.006 | 0.242 / 1.18 | 0.266 / 1.19 |
| 2 | **0.787**±0.008 | <0.2 | **0.789**±0.004 | 0.752±0.004 | 0.211 / 1.17 | 0.248 / 1.19 |
| 3 | **0.799**±0.008 | <0.2 | **0.784**±0.003 | 0.758±0.005 | 0.216 / 1.15 | 0.242 / 1.19 |
| 4 | **0.722**±0.003 | <0.2 | **0.850**±0.001 | 0.747±0.005 | 0.150 / 1.24 | 0.253 / 1.20 |
| **u=0.49 μm/sec** | | | | | | |
| Exp | Approach<br>β | Retraction<br>β | Approach<br>β | Retraction<br>β | Approach | Retraction |
| | Eq.(4) | Eq.(6) | GMM (eq.14) | GMM(eq.14) | $H_{GMM}$ / $H_{RA}$ | $H_{GMM}$ / $H_{RA}$ |
| 1 | **0.365**±0.002 | 0.161±5x10$^{-4}$ | **0.368**±0.023 | 0.317±0.022 | 0.632 / 1.22 | 0.683 / 1.21 |
| 2 | **0.416**±0.002 | 0.163±5x10$^{-4}$ | **0.415**±0.026 | 0.342±0.022 | 0.585 / 1.22 | 0.658 / 1.23 |
| 3 | **0.337**±0.002 | 0.162±6x10$^{-4}$ | **0.362**±0.019 | 0.332±0.020 | 0.638 / 1.23 | 0.668 / 1.22 |
| 4 | **0.331**±0.002 | 0.162±6x10$^{-4}$ | **0.388**±0.019 | 0.316±0.023 | 0.612 / 1.23 | 0.684 / 1.21 |

We have to notice that the linear fittings (eq.11) displayed in Figure S3 (lower panels) present a maximum error of 7%. For a viscoelastic environment where no other mechanism affects the time series the value of the structure function for q=1 corresponds to the Hurst exponent, see the points highlighted by a circle in the same figures. The upper panels of Figure S3 depicts the structure functions of the form $z(q)=Hq-C(q^2-q)$, where $H=H_{GMM}$ and C expresses intermittency.[5] This form of z(q) underlines the existence of a multiplicative mechanism of two competitive and independent sources shaping the overall response signal. The first one is assigned to the viscoelastic environment, and the second one is likely attributed to creep, or drift, or feedbacks from electronics. A quantification of them requires a more sophisticated treatment; a fractional Langevin equation with multiplicative noise as well with an additive fractional Gaussian noise could be a way of modeling it.[6] We leave this challenging task for future work.

**Section II**

Power spectrum scaling exponents obtained either directly by linear regression of (PSD), equation (15), ln(PSD)~a+ $\gamma$ ln(z), or indirectly through equation (16) of the main text. For the linear regression frequencies smaller than one fourth of the maximum frequency have been used. The results are listed in Table S2.

Table S2. Power spectrum scaling exponent for both approach/retreat phases

| | | Approach | | Retreat | |
|---|---|---|---|---|---|
| Exp | $v_0$ (μm/sec) | γ (eq.16) | γ (PSD) | γ (eq.16) | γ (PSD) |
| 1,2 | 0.0248 | -0.04±0.05,<br>0.22±0.028 | 0.47±0.19<br>0.33±0.17 | -0.62±0.062,<br>-0.44±0.03 | -0.60±0.25,<br>-0.60±0.21 |
| 3,4 | 0.0248 | 0.10±0.03,<br>0.20±0.032 | 0.11±0.15,<br>0.35±0.22 | -0.50±0.036,<br>-0.70±0.032 | -0.44±0.21<br>-0.52±0.18 |
| 5,6 | 0.0289 | 0.32±0.048,<br>0.26±0.044 | 0.60±0.21,<br>0.34±0.21 | -0.62±0.026,<br>-0.52±0.048 | -0.55±0.29,<br>-0.42±0.29 |
| 7,8 | 0.0289 | 0.24±0.054,<br>0.04±0.042 | 0.39±0.21,<br>0.35±0.27 | -0.46±0.07,<br>-0.60±0.048 | -0.48±0.26,<br>-0.42±0.30 |
| 9,10 | 0.0289 | 0.22±0.06,<br>0.04±0.04 | 0.22±0.21,<br>0.19±0.16 | -0.38±0.052,<br>-0.52±0.052 | -0.49±0.21,<br>-0.63±0.18 |
| 11,12 | 0.0289 | 0.18±0.054,<br>0.16±0.070 | 0.30±0.18,<br>0.33±0.23 | -0.52±0.062,<br>-0.34±0.034 | -0.51±0.30,<br>-0.23±0.33 |
| 13 | 0.0331 | 0.50±0.06 | 0.50±0.27 | -0.54±0.046 | -0.38±0.25 |
